# Transforming summary statistics from logistic regression to the liability scale: application to genetic and environmental risk scores

**DOI:** 10.1101/385740

**Authors:** Alexandra C. Gillett, Evangelos Vassos, Cathryn M. Lewis

**Author notes:** Corresponding Author Cathryn M. Lewis Institute of Psychiatry, Psychology & Neuroscience Social, Genetic & Developmental Psychiatry Centre King’s College London 16 De Crespigny Park Denmark Hill, London, SE5 8AF, UK.

## Abstract

**Abstract:** *Objective:* Stratified medicine requires models of disease risk incorporating genetic and environmental factors. These may combine estimates from different studies and models must be easily updatable when new estimates become available. The logit scale is often used in genetic and environmental association studies however the liability scale is used for polygenic risk scores and measures of heritability, but combining parameters across studies requires a common scale for the estimates.

*Methods:* We present equations to approximate the relationship between univariate effect size estimates on the logit scale and the liability scale, allowing model parameters to be translated between scales.

*Results:* These equations are used to build a risk score on the liability scale, using effect size estimates originally estimated on the logit scale. Such a score can then be used in a joint effects model to estimate the risk of disease, and this is demonstrated for schizophrenia using a polygenic risk score and environmental risk factors.

*Conclusion:* This straightforward method allows conversion of model parameters between the logit and liability scales, and may be a key tool to integrate risk estimates into a comprehensive risk model, particularly for joint models with environmental and genetic risk factors.

## 2 Introduction

Stratified medicine aims to improve health outcomes by developing targeted medical and public health interventions for individuals or subgroups of a population. A prerequisite of these strategies is the development of methodology to calculate the expected risk of developing a disease based on individual characteristics. Appropriate preventative strategies can then be applied, including enrolling participants in screening programmes or intervention therapies designed to modify lifestyle risk factors. Such strategies can improve disease outcomes for patients through earlier diagnosis and intervention and reduce the burden of disease in a population.

Statistical methods that appropriately model disease risk are therefore important. These models should be flexible, easily updated as new risk variables are found and pertinent across different populations [1]. To maximize the predictive power of these models, it is necessary to combine information on genetic and environmental risk factors, using existing risk estimates taken from studies with different designs and analysis methods. Crucially, a common scale across risks is required to combine the information of all disease associated variables. Here, scale refers to the function through which the risk variables relate to the risk of disease.

For environmental risk variables, the logit scale is most commonly used to model the relationship between the binary outcome of disease status and risk factor, in a logistic regression [2]. Similarly, logistic regression is used in genetic studies of disease, testing for association between a single nucleotide polymorphism (SNP) and disease status. However, some study designs perform analysis on the liability scale, with the summary measure of heritability being presented on this scale. Models of disease using the logit and the liability scale can be written as latent variable models, where it is assumed that the risk of disease is a function of a continuous, unobserved (latent) variable. The liability scale assumes that this latent variable is either normally distributed, or a mixture of normal distributions, while the logit scale assumes that this latent variable follows a logistic distribution, or a mixture of logistic distributions.

Both logit and liability scales are used in generating polygenic risk scores (PRSs). Within a discovery sample, SNP-by-SNP logistic regression is run for all SNPs from a genome-wide association study. A weighted sum of SNPs, weighting each SNP by log odds ratios (ORs), is then calculated with multiple p-value thresholds for SNP inclusion used to create multiple scores. The final PRS is selected in a target sample, where each score is regressed against disease outcome in turn and the best performing score, in terms of variance explained, is selected. One summary measure presented for the final PRS is the estimated proportion of the variability in liability to disease attributable to the PRS, which is calculated in the target sample and presented on the liability scale. Similarly, incorporating family history of disease into the joint effects model may also be easier using the liability scale. The disease outcome of multiple relatives can be used by defining the multivariate distribution for the latent variables underlining disease for all family members.

We therefore select the liability scale as the common scale for risk models combining existing risk estimates from the literature. Hence, equations to transform effect size estimates from the logit scale to the liability scale are required. Here, we present such equations which use the similarities between the cumulative distribution functions (CDFs) of the normal and the logistic distributions. This method requires the population prevalence of disease, the log OR for a risk factor and the frequency of the risk factor in the population. The resulting effect size estimates on the liability scale can be used within models of disease risk on the liability scale, which can combine risk information from multiple risk factors, obtained from different sources. We demonstrate these methods by estimating the risk of schizophrenia using a PRS and five environmental risk factors.

## 3 Methods

In this work we aim to transform the univariate log OR for a risk variable, which we also call the effect size on the logit scale, to the liability scale. To do this, we define models of disease on both scales. For the model on the logit scale we show that the resulting risk of disease is a function of the CDF for the logistic distribution. Similarly, we show that the risk of disease from the model on the liability scale is a function of the CDF for the standard normal distribution.

The approximate equivalence of the CDF for the standard normal distribution and the logistic distribution has long been noted [3-6]; an example of this can be seen in Figure 1. Having demonstrated that the conditional probability of disease from a model the logit scale can be written as a logistic CDF, and similarly for the liability scale as a standard normal CDF, we use this approximate equivalence to equate the disease risks from both models. We therefore generate an equation linking the log OR to the corresponding effect size on the liability scale. A joint effects model on the liability scale can then be built using these transformed effect size estimates, assuming the risk variables are independent and do not interact.

**Fig. 1.**
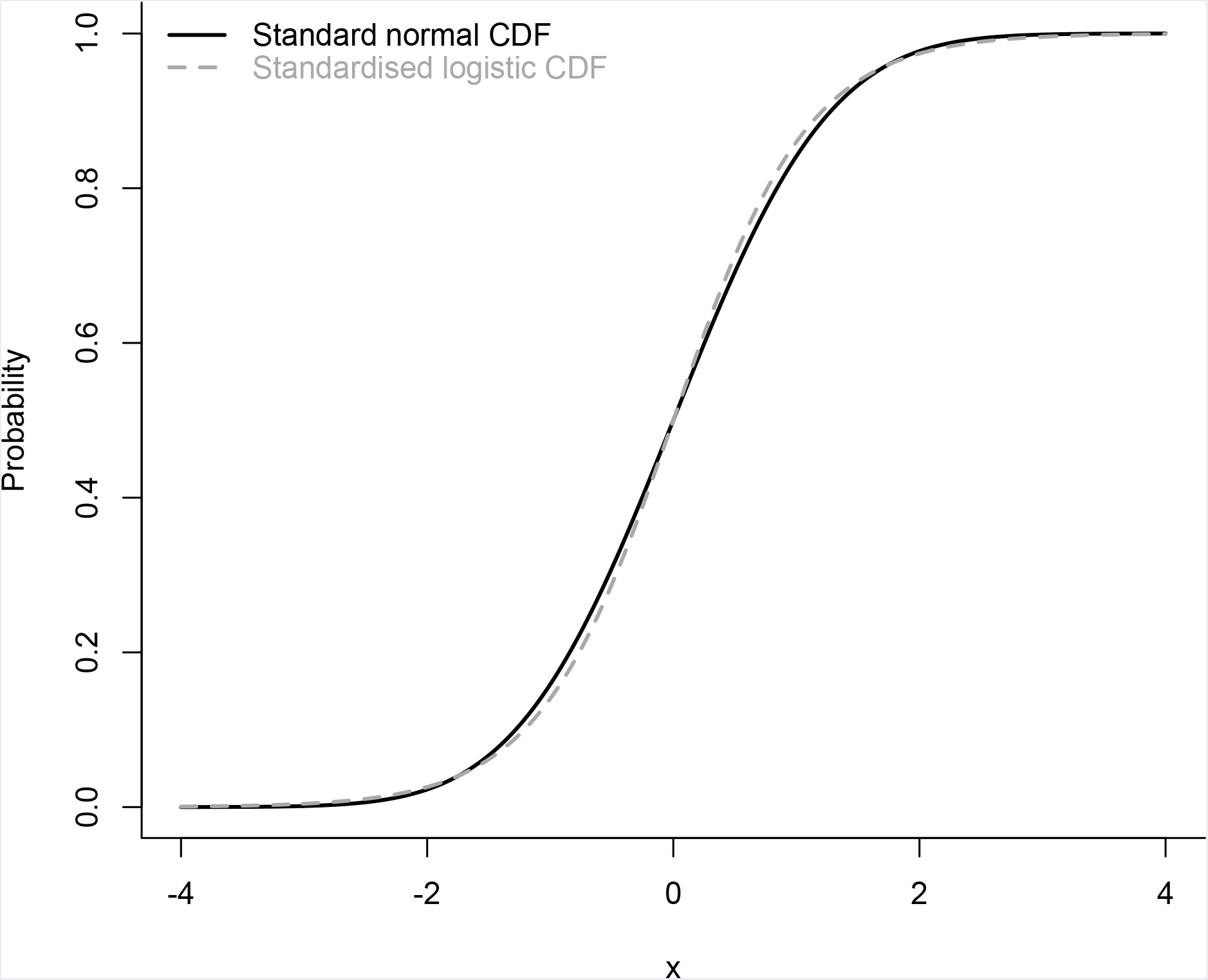
Plot of the normal and logistic cumulative distribution functions (CDF). The standard normal CDF is: 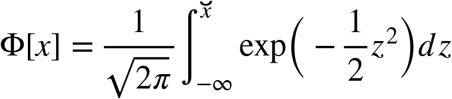. The standardised logistic CDF, where the mean is 0 and variance is 1 is given by: 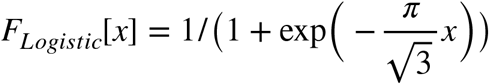.

To begin, we define the disease status variable, *Y* ~ *Binom*(1,*K*), where *Y* = 1 if an individual is affected with the disease of interest and 0 otherwise. *K* = *p*(*Y* = 1) is the population prevalence.

We assume that a risk factor, *X*, is observed. For simplicity, we assume that *X* is a discrete random variable with *η* + 1 categories, where: 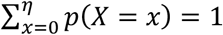.

We want to quantify the relationship between this risk factor and disease status using the liability scale. However, the relationship between a risk factor and disease outcome is typically estimated using logistic regression within a case-control study. We therefore transform the effect size estimates on the logit scale, the log ORs, to the liability scale. We now define the risk model on the logit and the liability scales, to understand what we are transforming from and to.

### 1. Defining the model on the logit scale

The statistical model relating the risk factor to disease outcome on the logit scale is the logistic regression model, and we therefore write the conditional risk of disease as:

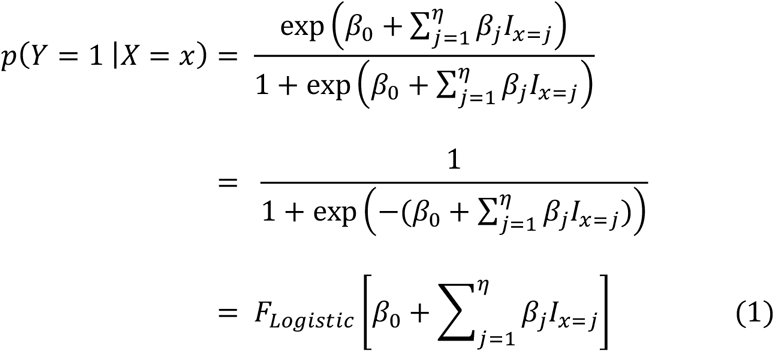

where:

- *β*_0_ is the intercept, corresponding to the log-odds of disease when {*X* = 0},
- *β_j_* is the change in the log-odds of disease from observing {*X* = *j*} compared to the reference category {*X* = 0}, and,
- *F_Logistic_*[*t*] is the CDF of the logistic distribution with 0 mean and variance equal to *π*^2^/3. Here we use a logistic distribution with zero mean and variance equal to *π*^2^/3 as this is the distribution used in the generation of effect size estimates from logistic regression outputted from standard statistical software, such as the glm function from the stats library within R.

### 2 Defining the model on the liability scale

We define a latent variable, *L*, known as the ‘liability to disease’, such that:

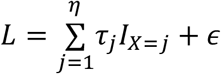

where:

- *τ_j_* is the change in liability to disease from observing {*X* = *j*} compared to the reference category {*X* = 0},
- *I*_*x*=*j*_ is a dummy variable which equals 1 if {*X* = *j*} and 0 otherwise (*j* = 1,…,*η*), and,
- *∊* ~ *N*(0,1 – *Var*[ξ(*X*)]), with 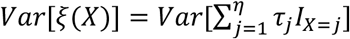.

*L* therefore has a unit variance (*Var*[*L*] = 1), as is standard in the definition of liability to disease. We note that the expected value of *L* here is not 0; instead we assume that *E*[*L*|*X* = 0] = *τ*_0_ = 0. This assumption is made for simplicity, and once model parameters *τ_j_* (*j* = 1,…,*η*) have been found, 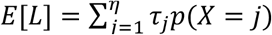 can be calculated and a mean centred liability to disease variable found if required.

Using the above definition of liability to disease, the probability of disease given the observed risk factor can be written as:

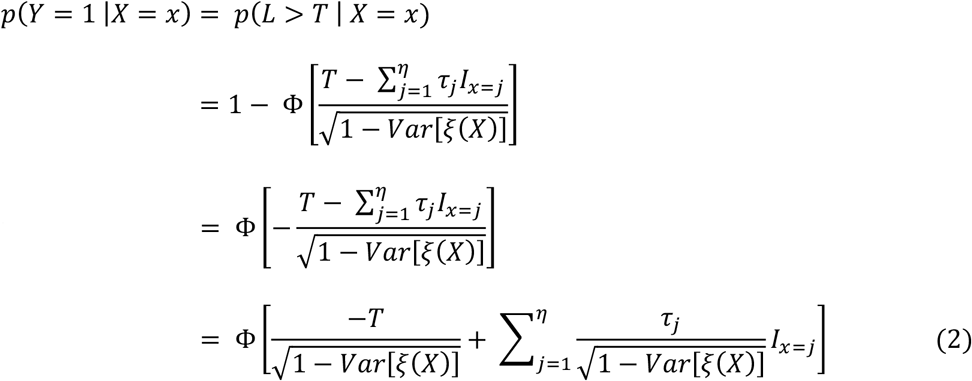

where 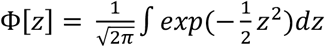 is the CDF of the standard normal distribution.

If the model relating disease status to the risk factor is defined on the liability scale, the risk of disease given the observed risk factor is a normal CDF. Similarly, if the model of disease is defined on the logit scale, the risk of disease given the observed risk factor is a logistic CDF.

The CDFs of the standard normal distribution and the unit variance standardised logistic distribution can be used to approximate one another [3]. Recall that output from logistic regression provides estimates of risk as a function of the logistic CDF with variance equal to *π*^2^/3, and not 1. However, we note that:

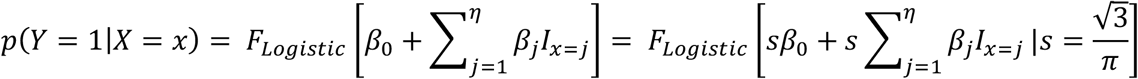

where:

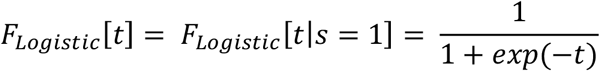

is, as before, the CDF corresponding to the logistic distribution with mean equal to 0 and variance equal to *π*^2^/3, and:

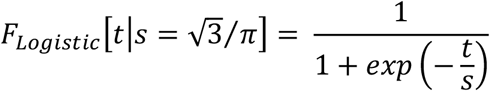

is the CDF corresponding to the standardised logistic distribution with mean equal to 0 and variance equal to 1. That is, if the logistic regression model were re-run using an error variable following the standardised logistic regression then the resulting coefficient estimates would equal the usual logistic regression coefficients multiplied by the scale parameter, *s*. Further, the estimated risk of disease would be the same regardless of whether the logistic regression used an error variable with a variance of *π*^2^/3 or 1.

Using the approximate equivalence of the standard normal CDF and the standardised logistic CDF, we expect the risk estimates from a model on the liability scale to be approximately the same as those on the logit scale. We therefore say:

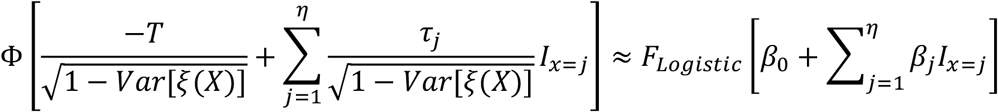

When {*X* = 0} is observed we can write the above approximation as:

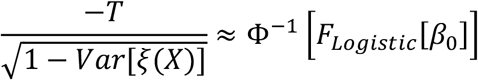

Then for all other observed values, {*X* = *j*};*j* = 1,…, *η*, we obtain:

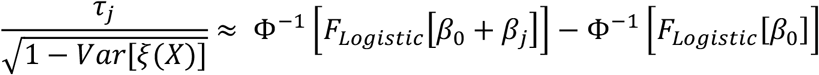

It can be shown that:

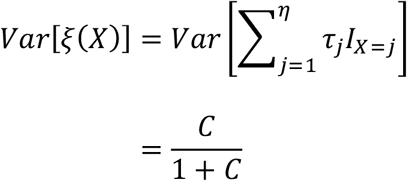

where *C* is a *calculable* constant defined as:

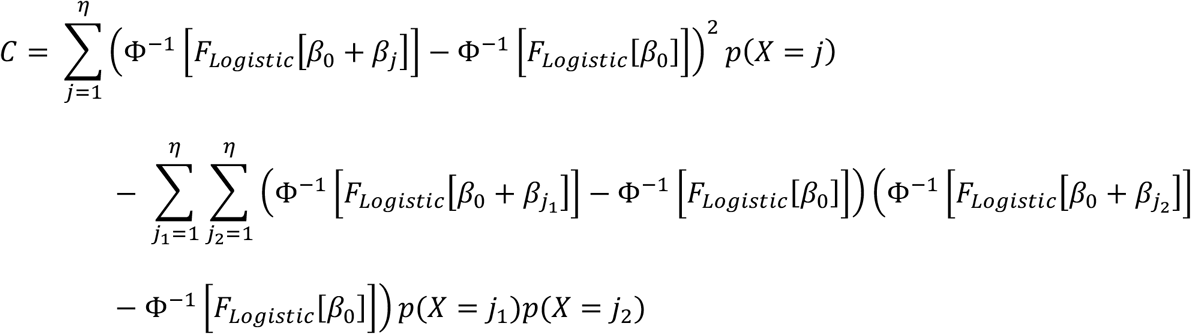

Details of this derivation are provided in the Supplementary Materials. This provides the following approximation for the effect size on the liability scale of the *j^th^* category for a risk factor:

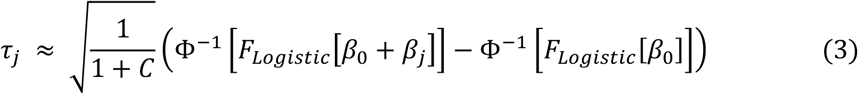

If we have model parameter estimates from a logistic regression model, including the intercept: {*β_j_*; *j* = 0,1,…, *η*}, then we can use the above approximation to estimate the corresponding effect size estimate on the liability scale.

We note that the intercept is required. The intercept is not typically reported. Additionally, this is the intercept assuming no ascertainment bias. It is common in case-control studies to over-sample cases. In logistic regression the over-sampling of cases impacts the intercept estimate, but not the effect size estimates. In either of these situations, the required population intercept, *β*_0_, can be calculated in the following manner. Using the law of total probability, and the logistic risk model, we can write:

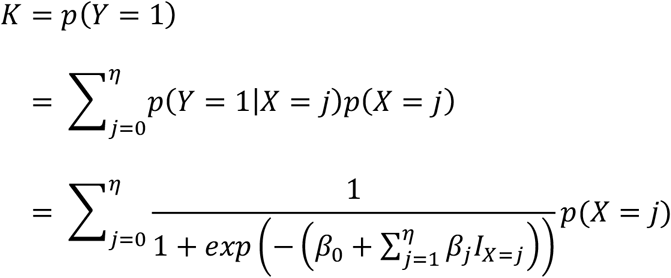

If we have estimates for the population prevalence, the log ORs for the risk factor of interest and the probability distribution function (PDF) for the risk factor, then the only unknown in the above equation is the required *β*_0_. This can then be calculated using numerical optimisation, such as by using the uniroot function within the stats R package. The approximation in Equation (3) can then be used to calculate *τ_j_*.

If we can assume that the effect of the risk factor, *X*, is additive on the liability and the logit scale, such that Equation (1) is:

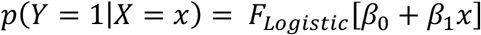

and Equation (2) is:

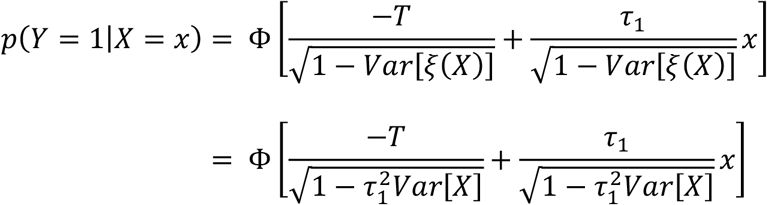

then the approximation to obtain the effect size estimate on the liability scale simplifies to:

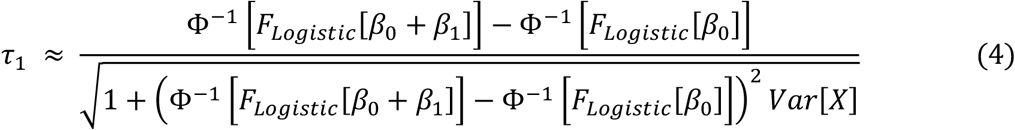

The approximation in Equation (4) is suitable for use when the risk factor under consideration is a SNP which is assumed to have an additive relationship with the liability to disease.

For any single risk factor, it may be reasonable to assume that its contribution to the total variability in liability to disease is very small. In such a case, we could assume that:

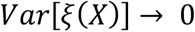

and the approximation becomes:

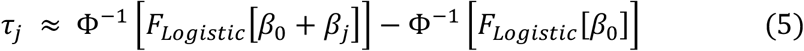

If *τ_j_* is small, as is the case for the effect size of SNPs in a PRS, or if the probability of being in a single category is large, for example for rare copy number variants, then:

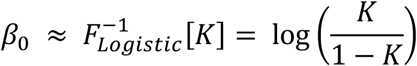

We can then achieve our aim of transforming univariate effect size estimates for a risk factor from the logit to the liability scale, by using this in Equation (5), giving:

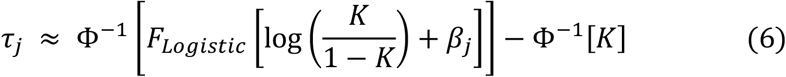

We now present an example applying this transformation. We estimate the risk of schizophrenia, integrating summary statistics for five environmental risk factors and a polygenic risk score (PRS) into a risk model on the liability scale.

## 4. Estimating the risk of schizophrenia

Schizophrenia is disabling mental health disorder, characterised by hallucinations, delusions or disordered thinking. The prevalence of schizophrenia is low (estimated lifetime prevalence of 1%), but the disorder has a huge personal and economic impact [7-9].

We aim to estimate an individual’s risk of schizophrenia using a polygenic risk score (PRS) and an environmental risk score (ERS).

The Psychiatric Genomics Consortium estimated that the proportion of variability in the liability to schizophrenia explained by the PRS was 0.07 [10]. We use this PRS summary measure, already on the liability scale, within our risk model and define the PRS random variable to be:

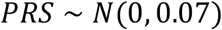

For the ERS, we combine information from meta-analyses of five environmental risk factors: cannabis usage, migrant status, urbanicity, paternal age and childhood adversity [11-15]. These meta-analyses present risk effect size estimates as ORs, except for migrant status, which uses relative risk. We opt to treat the relative risk as an OR here. In addition to the effect size estimates, the probability distribution function (PDF) for each risk factor is taken from its corresponding meta-analysis.

For each risk factor, the effect size estimate and distribution of the risk factor levels are extracted from the meta-analyses, and Equation (3) is used to convert each OR to their corresponding effect size on the liability scale (Table 1). The intercept for each risk factor on the logit scale is found using numerical optimization. R code for the transformation procedure is available from https://github.com/alexgillett/scale_transformation.

**Table 1.**
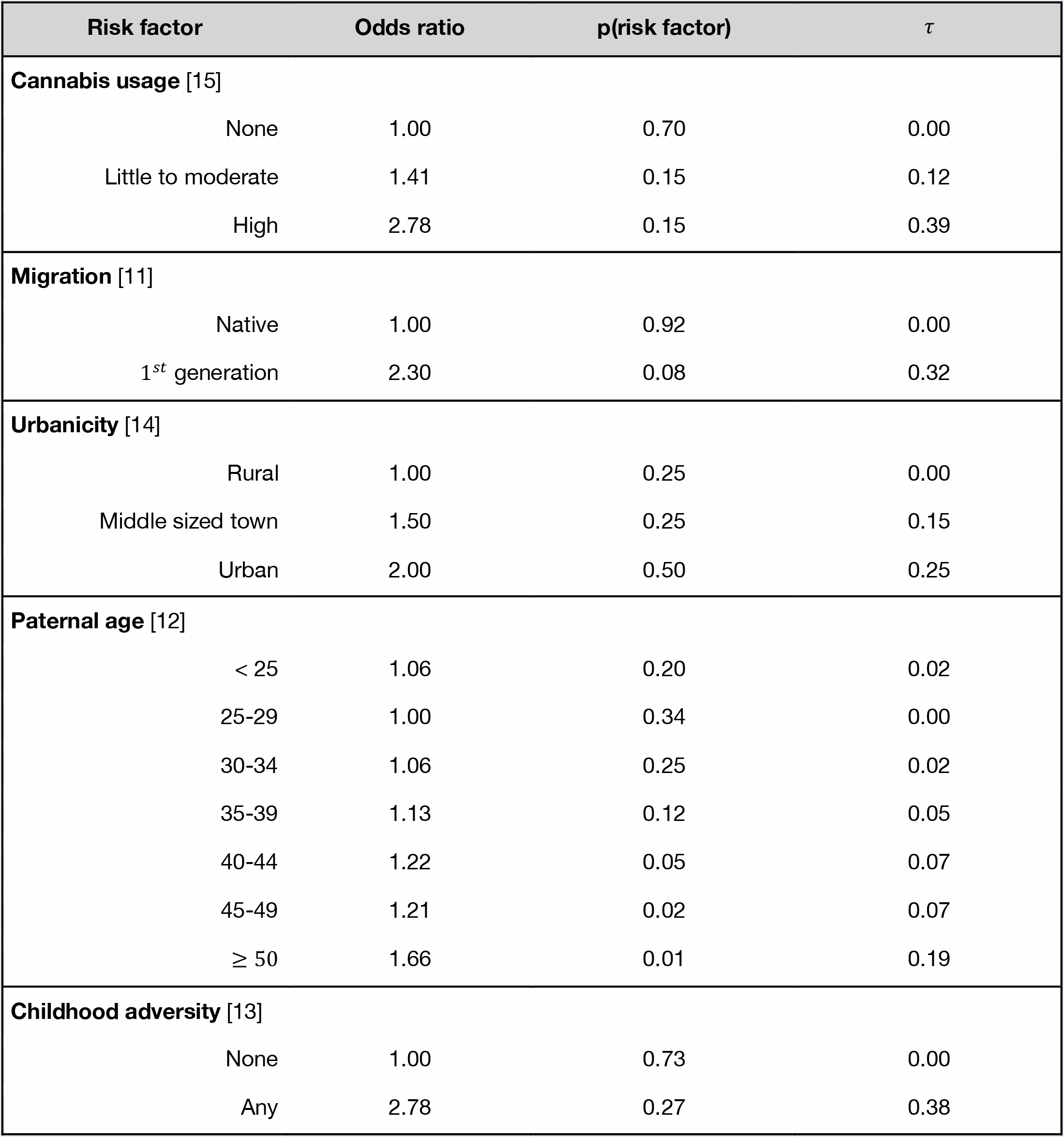
Environmental risk factors to be included in the schizophrenia environmental risk score. *τ* = effect size estimate on the liability scale, gained by transforming the odds ratio using Equation (3).

| Risk factor | Odds ratio | p(risk factor) | $\tau$ |
| --- | --- | --- | --- |
| **Cannabis usage** [15] |  |  |  |
| None | 1.00 | 0.70 | 0.00 |
| Little to moderate | 1.41 | 0.15 | 0.12 |
| High | 2.78 | 0.15 | 0.39 |
| **Migration** [11] |  |  |  |
| Native | 1.00 | 0.92 | 0.00 |
| 1<sup>st</sup> generation | 2.30 | 0.08 | 0.32 |
| **Urbanicity** [14] |  |  |  |
| Rural | 1.00 | 0.25 | 0.00 |
| Middle sized town | 1.50 | 0.25 | 0.15 |
| Urban | 2.00 | 0.50 | 0.25 |
| **Paternal age** [12] |  |  |  |
| < 25 | 1.06 | 0.20 | 0.02 |
| 25-29 | 1.00 | 0.34 | 0.00 |
| 30-34 | 1.06 | 0.25 | 0.02 |
| 35-39 | 1.13 | 0.12 | 0.05 |
| 40-44 | 1.22 | 0.05 | 0.07 |
| 45-49 | 1.21 | 0.02 | 0.07 |
| ≥ 50 | 1.66 | 0.01 | 0.19 |
| **Childhood adversity** [13] |  |  |  |
| None | 1.00 | 0.73 | 0.00 |
| Any | 2.78 | 0.27 | 0.38 |

Assuming that the risk factors to be included in the ERS are independent, and do not interact on the liability scale, we define the ERS component to be:

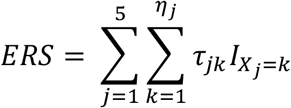

where *τ_jk_* is the change in liability to disease from observing the *K^th^* category for the *j^th^* risk factor (denoted *X_j_*), compared to observing the reference category for the *j^th^* risk factor; *K* = 1,…,*η_j_* and *j* = 1,…,5. The resulting schizophrenia ERS has 216 unique combinations, with values ranging from 0 to 1.53. A value of 0 corresponds to an individual in the lowest risk category for all five environmental risk factors. That is, someone who does not use cannabis, is native to the country in which they live, grew up in a rural setting, experienced no childhood adversity, and whose father was aged between 25 and 29 at conception. The ERS population average is:

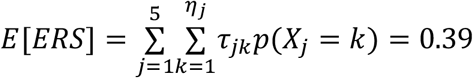

and the variance is:

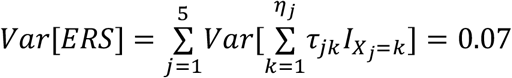

These five environmental risk factors therefore explain approximately the same proportion of the variability in liability to disease as the schizophrenia PRS. We now define the following joint effects model for schizophrenia as:

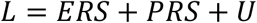

where *ERS* and *PRS* are as defined above, and *U* is the unmeasured liability to disease random variable, which is assumed to follow the normal distribution:

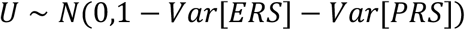

thereby ensuring that *Var*[*L*] = 1.

If the disease threshold parameter, *T*, was known then this risk model on liability scale provides the following equation to estimate the conditional risk of schizophrenia:

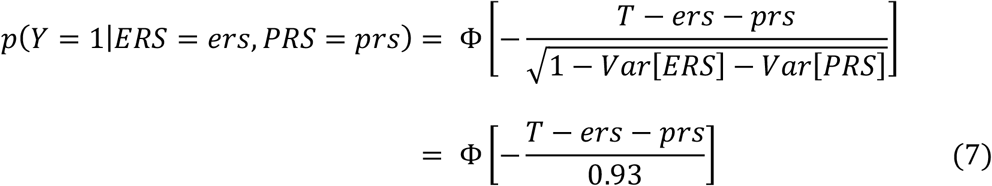

Assuming that the liability to disease, *L*, is normally distributed rather than a mixture of normal distributionS’ as is the case here, we can write:

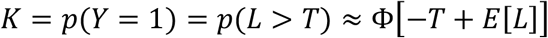

which provides the following approximation for *T*:

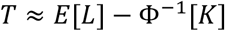

Here, *E*[*L*] = *E*[*ERS*], therefore giving an estimate of:

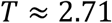

for the disease threshold. Using this in Equation (7) we can gain estimates of the risk of schizophrenia, given an observed ERS and PRS.

Estimates of risk from one risk score (PRS or ERS) can be calculated as:

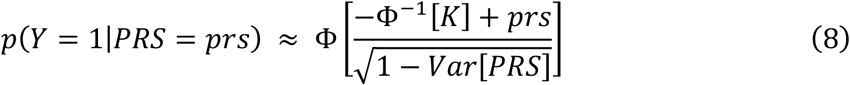

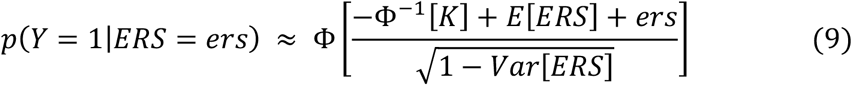

We now present estimates for the risk of schizophrenia from the PRS, the ERS and both scores, using Equations (7) - (9) (Figure 2). For ease of interpretation, the PRS is standardised to be from a standard normal distribution.

**Fig. 2.**
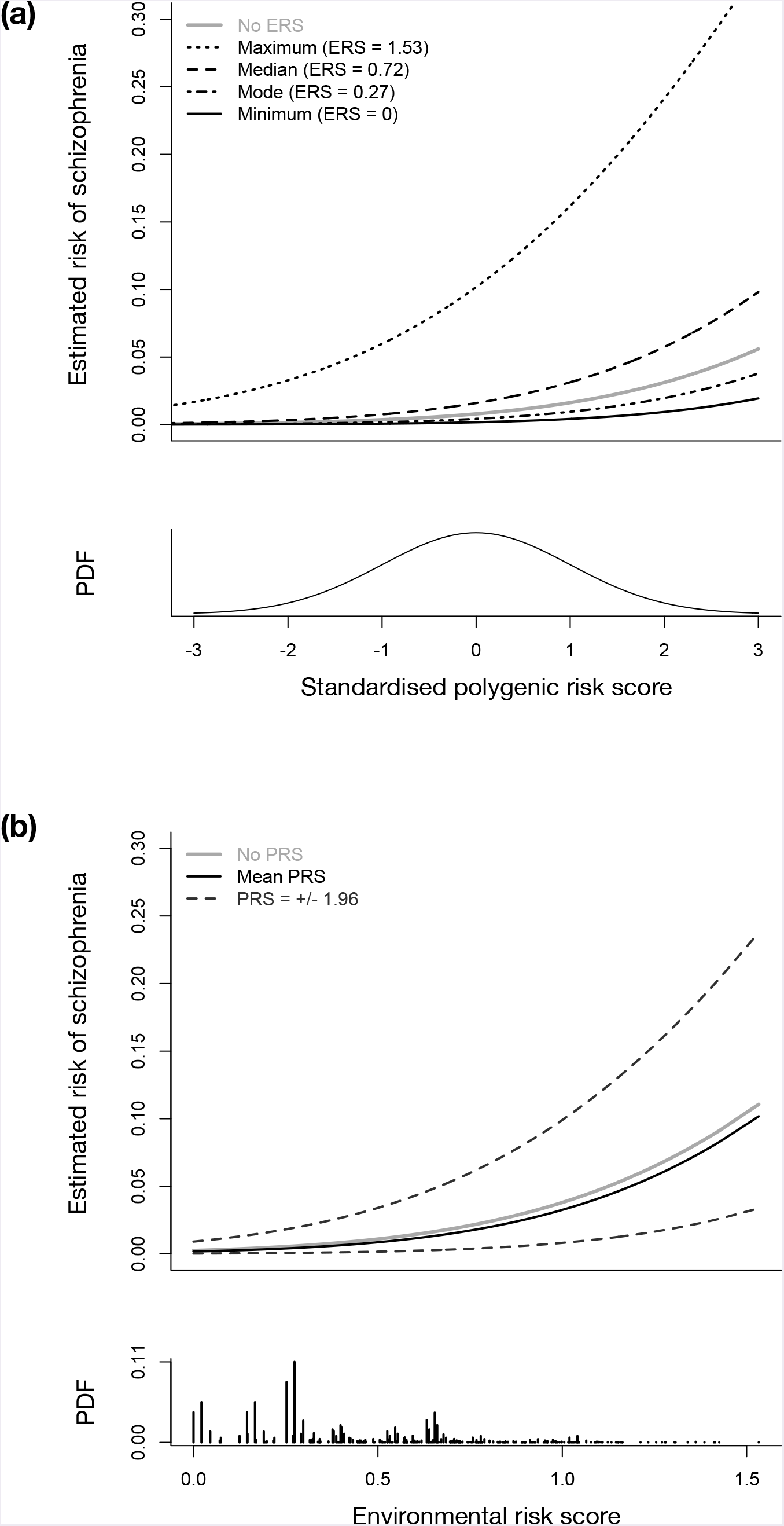
Estimated risk of schizophrenia against: (a) the standardised polygenic risk score (PRS), and, (b) the environmental risk score (ERS). Line types in plot: (a) refer to different ERS values including when no ERS is included in the model and, (b) refer to no PRS being included in the model and the 2.5^th^, 50^th^ and 97.5^th^ PRS percentiles. The probability density functions (PDFs) for: (a) the standardised PRS and (b) the ERS are also provided.

Under the PRS-only model (Figure 2a), 95% of individuals are expected to have a schizophrenia risk estimate between 0.0016 and 0.0304. An individual with a PRS > 95^th^ percentile has at least a 3-fold increased risk of schizophrenia, compared to the population prevalence of approximately 1%. However, only 2.5% of the population has a risk of this magnitude, and an individual with a PRS in the 95^th^ percentile still only has a low absolute risk of schizophrenia.

Under the ERS-only model (Figure 2b), the estimated for schizophrenia ranges between 0.0025 and 0.1107. An individual can therefore be as much as 11 times more likely to develop schizophrenia compared to the average individual in the population depending on their ERS. However, 95% of individuals are expected to have an estimated risk for schizophrenia between 0.0025 and 0.0267. Although a relatively high risk estimate can be achieved by considering the joint effects of environmental risk factors, it is rare to observe the ERS corresponding to this.

By using the joint effects model, with both PRS and ERS included, we can further refine the estimates of disease risk. For example, under the PRS-only model, an individual with an average PRS of 0 has an estimated risk of 0.0079. When the ERS is also included in the model, the estimated risk of schizophrenia ranges between 0.0017 and 0.1018. Including both PRS and ERS in the model therefore improves our ability to stratify individuals by risk. We do note that this upper risk estimate will be rarely observed; only 0.0006% of individuals expected to have a PRS equal to 0 and an ERS equal to 1.53. Indeed, under the joint effects model, 95% of individuals will have an estimated risk of disease between 0.0006 and 0.0458. As expected given the distribution of the risk scores in the population, and the impact of these risk scores shown under the single risk score models, most individuals have an estimated risk of disease low in absolute value.

However, this risk range from the joint risk model is wider than that found using the PRS and ERS only models, which suggests better calibrated risk estimates from the joint effects model [16]. Additionally, despite being a rare occurrence, for individuals with a high PRS and ERS, it is useful to understand the magnitude of their increased risk given our current knowledge.

## 5. Discussion

Motivated by building a joint effects model for disease risk on the liability scale using parameter estimates available in the literature, we have presented equations to transform effect size estimates from the logit to the liability scale. This method uses the approximate equivalence of the normal and logistic CDFs to define a relationship between the univariate effect size estimates for a risk factor on both scales. Assuming that the risk factors under consideration are independent and act additively, without interaction, on the liability scale then the resulting transformed effect size estimates can be used in a joint effects model to estimate the risk of disease. Such a model, which uses existing summary statistics in its parameterisation, is easily updated, and therefore may have utility in stratified medicine.

This approach was explored using schizophrenia where we constructed a joint effects model for risk using a polygenic risk score (PRS) and five environmental risk factors. Using schizophrenia associated risk variables from different studies, we created an environmental risk score (ERS) on the liability scale from effect size estimates on the logit scale, and fully parameterised the liability model by approximating the disease threshold parameter. We demonstrated the potential improved ability of the joint effects model to stratify individuals by risk compared to the single risk score models based on only PRS or ERS. Individually, the PRS and ERS each explain 7% of the variability in liability to disease. Combined, and assuming independence, they explain 14%. Further improvements in risk stratification for schizophrenia will come as knowledge of risk variables expands, and this model is updated.

In this work we have presented equations to transform univariate effect size estimates from the logit scale to the liability scale. To incorporate the transformed effect size estimates from multiple risk variables into a joint effects model for disease risk on the liability scale we must therefore assume that the risk variables are independent and do not interact on the liability scale. Future work will investigate how to relax these assumptions, however the simplest way to do this is likely to use effect size estimates on the logit scale from studies considering the joint effects of correlated and interacting risk variables.

It is possible to build joint effects models on other risk scales, for example the logistic scale. The transformation equations presented here can, of course, be rearranged such that we transform effect size estimates on the liability scale to the logistic scale. However, as noted in the Introduction, there are benefits to using the liability scale when including the family history of disease in the joint effects model, with a stronger body of research in multivariate normal (MVN) distribution compared to the multivariate logistic (MVL) distribution. Indeed, since the MVL distribution is part of the extreme value distribution family, it has multiple potential definitions [17]. It is therefore simpler to use the liability scale, rather than the logit scale, in this circumstance and equations for risk estimation incorporating a PRS and family history have been derived using the liability model [18]. Although we do not consider family history here, we acknowledge that family history is important for stratifying individuals by risk. The methods for scale transformation presented here could be used in conjunction with a MVN approach to incorporate family history, allowing disease risk models to be constructed using environmental, genetic and family history risk profiles.

In addition to the logistic and liability models for disease risk, time-to-event models, such as the Cox proportional hazards model, may be used. Such models have the advantage of incorporating age at onset into the model, and therefore work with incidence rather than prevalence. Future work would be to investigate the parameterisation of such incidence models using commonly available summary statistics. The motivation for the scale transformation equations was to estimate disease risk using a PRS and environmental risk factors on the liability scale. Other uses for the transformation equations would be to transform SNP effect size estimates to be used in the estimation of SNP heritability. In summary, the equations developed here provide a straightforward method to convert model parameters from the logit scale to the liability scale (or *vice versa*), and may be a key tool to integrate risk estimates from published studies into a comprehensive risk model for disease risk, particularly allowing joint models across environmental and genetic risk factors to be constructed.

## 6 Statements

### 6.1 Statement of Ethics

The authors have no ethical conflicts to disclose.

### 6.4. Author Contributions

Alexandra C. Gillett derived mathematical equations, wrote R-code, created figures, tables and supplementary materials, and contributed to the writing of this manuscript.

Evangelos Vassos provided expert knowledge on schizophrenia, including variables for inclusion in the environmental risk score and their odds ratios, and contributed to the writing of this manuscript.

Cathryn M. Lewis provided expert knowledge in statistical genetics and schizophrenia, and contributed to the writing of this manuscript.

